# Bacteriophage-derived endolysins restore antibiotic susceptibility in penicillin- and erythromycin-resistant *Streptococcus pneumoniae* infections

**DOI:** 10.1101/2025.03.18.644011

**Authors:** Niels Vander Elst, Lisa Knörr, Kristine Farmen, Federico Iovino

## Abstract

*Streptococcus pneumoniae*, the pneumococcus, is a cause of major illness globally. Invasive pneumococcal disease (IPD) is characterized by pneumococci invading blood (bacteremia), lungs (pneumonia), or brain and cerebrospinal fluid (meningitis). Meningitis remains an important global health concern because half of the survivors experience long-term neurological damage. The antibiotics commonly used to treat pneumococcal infections are β-lactams and macrolides, however, *S. pneumoniae* is nowadays often resistant to one or several antibiotics, therefore novel antimicrobials are needed. Here, we found that the bacteriophage-derived Cpl-1 endolysin showed consistent antibacterial activity against β-lactam– and macrolide-resistant pneumococcal clinical strains grown in human blood and human cerebrospinal fluid. Exploiting synergistic and additive mechanisms, supplementation of cpl-1 to either penicillin or erythromycin rescued human neuronal cells from the cytotoxicity of antibiotic-resistant pneumococcal infections. Finally, systemic administration of cpl-1 supplemented to penicillin in mice infected with penicillin-resistant pneumococci successfully reduced bacteremia, and, thanks to the efficient penetration across the blood-brain barrier, abolished bacterial load in the brain, resulting in increased (89%) survival accompanied by an asymptomatic course of infection. These findings strongly indicate that cpl-1 can restore antibiotic sensitivity against β-lactam– and macrolide-resistant *S. pneumoniae*, representing a fundamental adjunct therapy to standard-of-care antibiotics against multidrug-resistant IPD.

## Introduction

*Streptococcus pneumoniae*, the pneumococcus, is a cause of major illness globally, and, because of its high incidence of resistance towards several antibiotics, was listed among the bacterial priority pathogens in 2024 by the World Health Organization (WHO) and classified as a “serious” threat by the Centers for Disease Control and Prevention^1,2^. The most severe manifestation of infection is invasive pneumococcal disease (IPD), characterized by pneumococci invading tissues and organs, such as blood, lungs, brain and cerebrospinal fluid leading to bacteremia, pneumonia and meningitis, respectively^3^. Bacterial meningitis, in which *S. pneumoniae* is the major cause globally^4^, is a life-threatening inflammation of the meninges caused by a bacterial infection of the brain and remains a major health burden globally. Up to 50% of survivors frequently suffer from permanent neurological disabilities due to neuronal damage caused by the infection^5,6^. Bacterial meningitis during childhood causes a significant higher risk to develop long-term neurological disorders later in adult lives, such as motor and cognitive impairment, visual and hearing loss, behavioral disturbances, and structural damage to the head^7^. Vaccination is in place as a preventive strategy for IPD, but the emergence of serotypes that circumvent coverage by the pneumococcal vaccines, known as serotype replacement, indicates the continuous need for an effective strategy against this disease^8^. As such, antibiotics currently constitute the most effective way to treat IPD, including pneumococcal meningitis^9^.

The current standard-of-care antibiotic treatment consists of either β-lactams, mainly penicillin, and macrolides, or in case of β-lactam-resistance, mainly erythromycin; alternatively, vancomycin is also used in the case of multidrug-resistance.

In addition, the emergence of antimicrobial resistant strains has become alarming, with specific concerns for increasing resistance against β-lactams and macrolides for pneumococcal infections^10–12^. To emphasize the urge for novel antimicrobials needed, the WHO announced that antimicrobial resistance (AMR) will become a major health issue due to the increasing levels reported, also for pneumococcal meningitis^10^. Simultaneously, the number of antimicrobials under development is insufficient to tackle the challenge of increasing emergence and spread of AMR^13^. To tackle AMR emergence, endolysins derived from bacteriophages have gained increasing attention in veterinary medicine due to (i) their rapid method of action, and (ii) high specificity for the target bacteria^14–18^. Endolysins are peptidoglycan hydrolases that degrade the Gram-positive cell wall resulting in osmotic lysis of the bacteria^16–18^. Endolysins are one of the major classes of novel antimicrobials under development according to the WHO^13^, with cpl-1 (a dimer) and cpl-7s (a monomer) being previously characterized as effective endolysins against the pneumococcus^19–26^. Remarkably, the knowledge on how endolysins can be beneficial in case of multidrug-resistant bacterial infectious disease, in particular meningitis, is still scarce^25,26^.

Through *in vitro* infection experiments using human blood, cerebrospinal fluid and neuronal cells, and *in vivo* through our established bacteremia-derived meningitis mouse model, we provided evidence that cpl-1 endolysin restores antibiotic susceptibility in β-lactam– and macrolide-resistant pneumococcal infections, preventing neuronal cell death, dramatically reducing bacterial load in the blood, in the heart and, importantly, completely abolishing bacterial presence in the brain thanks to cpl-1’s efficient capability to cross the blood-brain barrier. Furthermore, endolysin treatment was also capable of suppressing inflammation, therefore promoting the maintenance of a safe environment for preserving normal neuronal activity. Endolysins should therefore be considered as an essential adjunctive to the current standard-of-care antibiotics to fight multidrug-resistant bacterial invasive disease.

## Materials and Methods

### Plasmid construction and *de novo* synthesis

The cpl-1 and cpl-7s coding sequences were codon optimized for expression in *E. coli* and chemically synthesized by Twist Bioscience (California, USA). A C-terminal hexahistidine-tag was included for purification purposes. All constructs were cloned into a pET28a(+) vector. Benchling (Biology software, 2024) was used for DNA sequence analysis and manipulations.

### Plasmid transformation, protein expression and purification

For *in vitro* experiments, chemocompetent *E. coli* BL21 (DE3) (New England Biolabs) were transformed with 25µg (5.0 ng/μL) plasmid via heat shock during 45 sec at 42°C after a 30 min incubation on melting ice. The transformed cells were then incubated for 1 h at 37°C in super optimal broth after a 5 min recovery on melting ice. *E. coli* cells were subsequently plated on Luria-Bertani (LB) agar (Oxoid) with the addition of 100 µg/mL kanamycin sulfate (Gibco). One colony was picked, transferred to a culture tube with 5mL LB and 100 ug/mL kanamycin sulfate, and incubated at 37°C and 200 rotations per minute (rpm) during 18 h. Next, overnight cultures were diluted 1:100 in LB broth supplemented with 100 µg/mL kanamycin sulfate and grown in Erlenmeyer flasks at 37°C and 200 rpm. When the OD_620nm_ reached 0.6 to 0.8, protein expression was induced by the addition of 0.5 mM isopropyl β-D-1-thiogalactopyranoside (Fisher Scientific) while shaking the Erlenmeyer flasks at 100 rpm at 21°C during 18 h. Subsequently, *E. coli* was pelleted and resuspended in 10 mL B-PER (Fisher Scientific) with the addition of 10 µL of 2500 U/mL DNaseI (Fisher Scientific). Bacterial suspension was then incubated on melting ice for 15 min on a shaker at 40 rpm. Next, centrifugation (4000 *g*; 20 min) was performed to pellet cellular debris, and the lysate was applied to a His Gravitrap column (Cytiva). This column was washed with 4.0 mL of: (i) lysis buffer (phosphate-buffered saline (PBS) with 10 mM imidazole (Carl Roth)), (ii) wash buffer (PBS with 50 mM imidazole and 1.0 M NaCl; pH 7.4) and (iii) eluted from the column in storage buffer (PBS with 500 mM imidazole, 0.5 M NaCl and 10% glycerol; pH 7.4) and kept at –80°C. After confirming correct protein expression and purification, large-scale protein production was conducted by the Protein Science Facility (PSF) at Karolinska Institutet. This involved a two-step purification process, being immobilized metal affinity chromatography followed by size-exclusion, performed on an Äkta Pure system (Cytiva). Protein sizes were verified by PSF using MALDI-TOF mass spectrometry.

For *in vivo* experiments, lipopolysaccharide (LPS)-deficient electrocompetent ClearColi BL21 (DE3) (Biosearch Technologies) were transformed with plasmid DNA by electroporation following the manufacturer’s protocol. Protein expression and purification were carried out similarly to the *in vitro* experiments, with an additional step to remove residual LPS. Pierce™ high-capacity endotoxin removal spin columns (Fisher Scientific) were used, resulting in protein with LPS levels below 0.1 EU/mL (essential for intravenous injection in mice), as measured by the Pierce™ chromogenic endotoxin quant kit (Fisher Scientific). The day before the experiment, a buffer exchange was performed to replace elution buffer with PBS using a Pierce™ protein concentrator with a 10.0 kDa molecular weight cut-off (Fisher Scientific).

### Pneumococcal culture conditions and whole genome sequencing

*S. pneumoniae* isolates were cultured either on blood agar plates (Karolinska Hospital, Sweden) or in Todd-Hewitt broth supplemented with 0.5% yeast extract (THY) (Karolinska Hospital, Sweden) and incubated at 37 °C in a 5% CO₂ atmosphere. For whole-genome sequencing, a single colony was selected from a plate and grown in THY for 18 h at 37 °C with 5% CO₂.

Genomic DNA was then extracted using the PureLink™ Genomic DNA Mini Kit (Fisher Scientific) following the manufacturer’s instructions. DNA of sufficient quality was sent to Plasmidsaurus (Oregon, USA) for sequencing and genome assembly. The resulting sequencing data, in FASTA format, were submitted to the PubMLST database (https://pubmlst.org/organisms/streptococcus-pneumoniae) (accessed on 30 September 2024) to confirm the bacterial species and identify allelic matches. Each *S. pneumoniae* isolate was characterized by its allelic profile, defined by allele numbers at seven loci: *aroE*, *gdh*, *gki*, *recP*, *spi*, *xpt*, and *ddl*. Based on the combination of these alleles, the sequence type (ST) was determined.

### Determination of the minimal inhibitory concentration and checkerboard assays

Minimum inhibitory concentrations (MICs) were determined according to EUCAST standards^27^. A bacterial inoculum of 5 × 10^5^ CFU/mL was prepared in Mueller-Hinton Fastidious (MH-F) broth (Karolinska Hospital, Sweden) supplemented with 5% lysed horse blood and 20 mg/L β-NAD (Svenska LABFAB). Plates were incubated for 18 h at 37°C, and bacterial growth was assessed by measuring the OD_620nm_ values using a spectrophotometer. The MIC was defined as the lowest concentration of antimicrobial that inhibited bacterial growth. The antimicrobials tested included benzylpenicillin (Meda), ceftriaxone (Navamedic), and vancomycin (MP Biomedicals), each with a clinical resistance breakpoint of 2.0 µg/mL. Erythromycin (Fisher Scientific) was tested with a breakpoint of 0.25 µg/mL. Additionally, the endolysins cpl-1 and cpl-7s were tested, although breakpoints for these agents have not yet been defined.

Checkerboard assays were conducted to evaluate potential synergy, additive, or antagonistic effects between endolysin cpl-1 and either penicillin or erythromycin. In brief, both antimicrobials were tested individually and in combination at varying concentrations to calculate the fractional inhibitory concentration (FIC). The FIC index was used to interpret the interaction, with FIC values indicating synergy (FIC ≤ 0.5), additive effects (FIC 0.5 – 4.0), or antagonism (FIC > 4.0). More detailed information on the checkerboard assay can be consulted in ^28^.

### Evaluation of the endolysin’s lytic and antibacterial activity

The peptidoglycan hydrolyzing and bactericidal activity of the endolysin was quantified using turbidity reduction (TRA) and time-kill assays (TKA) detailed in ^17^. For the TRA, a mid-log phase bacterial culture was pelleted (4000 *g*, 10 min), washed with PBS, resuspended, and diluted 1:1 in PBS to an OD_620nm_ of approximately 1.0. Then, 100 µL of the bacterial suspension was mixed with an equal volume of 5.0 µM endolysin in PBS (resulting in a final concentration of 2.5 µM) or with PBS alone as a negative control. OD_620nm_ was measured kinetically every 30 sec for 1 h at 37°C using a plate reader, shaking between readings. The enzymatic activity was calculated as (ΔOD_620nm_/min)/µM, standardized for endolysins as previously described in^29^. The TRA was also conducted in pooled human cerebrospinal fluid (Medix Biochemica, USA) by replacing PBS with cerebrospinal fluid in the assay. For the TKA, blood obtained from healthy volunteers (Karolinska sjukhuset, Sweden) was spiked with mid-log phase pneumococci and subsequently treated with a final concentration of 5.0 µM cpl-1 endolysin or PBS as a negative control. After 2 h of incubation at 37°C, serial dilutions were made and plated on blood agar plates to determine the remaining CFU/mL after incubation during 18 h at 37°C and 5% CO_2_.

### Culture, differentiation and infection of SH-SY5Y neuronal-like cells

Human SH-SY5Y neuroblastoma cells (ATCC) were cultured as previously described by our group. Briefly, cells were seeded in 12-well plates at a density of 100.000 cells/mL, differentiated for 10 to 14 days using differentiation media consisting of EMEM (ATCC) and F12 (Gibco) in 1:1 ratio, 5% fetal bovine serum (FBS) (Gibco), 1% penicillin/streptomycin (Gibco), and 10 μM retinoic acid (RA) (Bio-Techne) and incubated at 37°C with 5% CO_2_. The day before infection, neuronal-like cells were put into differentiation media without antibiotics as previously described^30^ (EMEM and F12 in 1:1 ratio, 2.5% FBS, 10 μM retinoic acid). The day of the experiment, the medium was discarded, cells were washed twice with PBS and pneumococci were resuspended in differentiation media without antibiotics and added to the neurons at a multiplicity of infection (MOI) of 10. The plates were centrifuged at 50 *g* for 5 minutes, and pneumococci were allowed to interact with the cells for 90 min before treatment was initiated and incubated at 37°C with 5% CO_2_. This treatment was added at a 1:1000 dilution rate.

### Animal experiments and tissue collection

Five-to six-week-old male C57BL/6J mice (JAX™, Charles River) were housed under standard conditions, with a 13:11 light/dark cycle and *ad libitum* access to food and water, in accordance with Swedish legislation (Svenska jordbruksverket), and ethical approval (permit number 18965–2021). A bacteremia-derived meningitis mouse model was employed as previously described by our group ^31^. Briefly, each mouse received an intravenous injection via the tail vein of 10^8^ CFU of *S. pneumoniae* isolate AMR 12,116 suspended in PBS. Two hours post-infection, mice were administered intravenously (tail vein) with 100 µL PBS (placebo), 100 mg/kg benzylpenicillin, 40 mg/kg of endolysin cpl-1, or a combination of both. The mice were monitored every 3 h for clinical signs, which were scored based on Karolinska Institutet’s veterinary protocol for assessing rodent welfare. Once an individual score or cumulative score reached 0.4 (considering 0.0 the clinical score of a healthy mouse), the animals were anesthetized using isoflurane, 100 µL of blood was obtained and the mouse perfused via the left ventricle with PBS, spleen, heart and brain were collected. For the assessment of BBB penetration of cpl-1 endolysin, mice were intravenously injected with 40 mg/kg of cpl-1 in 100 µL PBS and euthanized after one hour. Next, harvested organs were placed onto a 40 µm cell strainer (Corning Falcon) with the addition of 1 mL sterile PBS and crushed through the strainer with the plunger of a sterile syringe. The homogenates obtained were serially diluted, plated on blood agar plates, and incubated for 18h at 37°C and 5% CO_2_. The CFUs were counted and calculated back to CFU/mL the subsequent day. The remaining homogenates were immediately prepared for storage by adding 1% proteinase inhibitor cocktail (Fisher Scientific), allowed to rest at 4°C for 15 minutes, and stored at –20°C for further analyses.

### ELISA for IL-6 on mouse tissue

Upon thawing homogenates on ice, lysates were obtained after centrifugation (13,000 *g*, 3 min.), and protein concentration thereof determined on 1:10 dilutions by bicinchoninic acid assay (BCA) (Fisher Scientific). Standardization was done by dilution in PBS to equal protein concentrations. Quantification of IL-6 was performed by ELISA (Bio-Techne) according to the manufacturer’s protocol, loading 100 µg of protein into each well, in duplicate.

### Immunofluorescent staining of mouse tissue, microscopy analysis and signal quantification

Harvested brains were placed in 4% paraformaldehyde (Histolab) and post-fixed overnight at 4°C, placed in a 30% sucrose solution for a minimum of 48 h, and subsequently cut into coronal sections of 30 μm on a microtome (Leica SM2000). These sections were frozen in a cryoprotective solution (30% sucrose and 2% DMSO in PBS) and stored at –20°C until further processing. Prior to staining, the sections were washed in PBS, blocked with 5% goat serum (Gibco) and permeabilized with 0.3% Triton-X-100 (Sigma) for 1 h at room temperature, and subsequently stained for 2 h at room temperature with Alexa Fluor 647 conjugated anti-His (1:200, Abcam, AB237337-1001), for the detection of cpl-1 endolysin, and Alexa Fluor 488 conjugated lectin (1:250, VectorLabs) for the detection of the BBB vasculature^31^. Next, sections were washed in PBS and mounted with ProLong diamond mountant (Invitrogen). Images were obtained taking a z-stack of 20 μm of 1 μm images at 20× magnification using a Mica Microhub Imaging System (Leica) with settings kept constant during acquisition.

### Statistical analysis

Data were analyzed using GraphPad Prism (version 10.1.2) to calculate p values and determine statistically significant differences (p < 0.05). Data were analyzed as normally distributed datasets unless otherwise specified, confirmed by performing an Anderson–Darling test (n > 6) or by quantile–quantile plotting the residuals (n ≤ 6). Regarding pro-inflammatory cytokines, outlier values were detected using the ROUT method with Q set to 5% and removed from the dataset. Two groups were compared with two-tailed, (un)paired t tests, and multiple groups with analysis of the variance (ANOVA) and a Bonferroni post hoc test. Kaplan-Meier curve was analyzed by means of a log-rank (Mantel-Cox) test.

## Results

### The cpl-1 and cpl-7s endolysins exhibit antibacterial activity against antibiotic-resistant *S. pneumoniae* clinical isolates

A panel of clinically relevant pneumococcal isolates, collected from patients with IPD (bacteremia and sepsis) or meningitis, was assembled (**Table 1**). This panel included the reference laboratory strains *S. pneumoniae* TIGR4 (serotype 4) and D39 (serotype 2), along with 12 additional clinical isolates representing in total 11 serotypes. Notably, the panel featured serotypes not included in the currently available pneumococcal conjugated vaccines, such as 15A, 16F, 35A and 35B^32^. Each isolate was tested for antibiotic resistance against standard-of-care treatment according to EUCAST guidelines^27^. This analysis identified one penicillin-resistant serotype 35B isolate, with a MIC of 5.0 µg/mL, one erythromycin-resistant serotype 6A isolate, with a MIC of 4.0 µg/mL, and two isolates belonging to serotypes 23F and 35B resistant to both penicillin, with MIC of, respectively, 6.0 and 2.5 µg/mL, and erythromycin, both with MIC of 2.0 µg/mL.

**Table 1.** Pneumococcal isolates of this study. The reference laboratory strains D39 and TIGR4 as well as 12 clinical isolates derived from either invasive pneumococcal disease (IPD) or meningitis patients were tested for resistance against penicillin (MIC > 2.0 µg/mL) as well as erythromycin (MIC > 0.25 µg/mL). * Indicates a MIC above the EUCAST clinical breakpoint for resistance, whereas n.d. means not determined.

| Isolate | Serotype | Origin | MIC penicillin (µg/mL) | MIC erythromycin (µg/mL) | Piliated |
| --- | --- | --- | --- | --- | --- |
| D39 | 2 | IPD | 0.08 | 0.06 | no |
| TIGR4 | 4 | Meningitis | 0.08 | 0.03 | yes |
| AH 4,393 | 23F | IPD | 0.08 | 0.01 | n.d. |
| AH 16,031 | 35A | IPD | 1.25 | 0.03 | n.d. |
| AH 18,120 | 6A | IPD | 0.60 | 0.01 | n.d. |
| AMR 6A | 6A | Meningitis | 0.15 | 4.00 * | yes |
| AMR 12,116 | 35B | IPD | 2.50 * | 2.00 * | yes |
| AMR 19,000 | 35B | IPD | 5.00 * | 0.06 | yes |
| AMR 68,715 | 23F | Meningitis | 6.00 * | 2.00 * | yes |
| CCUG 35,561 | 18C | Meningitis | 0.08 | 0.06 | n.d. |
| CCUG 32,535 | 22F | Meningitis | 0.08 | 0.03 | n.d. |
| CCUG 35,268 | 9V | Meningitis | 0.08 | 0.06 | n.d. |
| FI 15A | 15A | Meningitis | 0.08 | 0.01 | yes |
| FI 16F | 16F | Meningitis | 0.08 | 0.03 | yes |

All pneumococcal clinical isolates were challenged with the cpl-1 and cpl-7s endolysins, and, as a result, all isolates were sensitive to the action of both endolysins (**Figure 1A**, **Supplementary Figure S1**). More specifically, a biochemical characterization was done through TRA which revealed that the cpl-1 compared to the cpl-7s endolysin hydrolyzed the peptidoglycan of these isolates significantly faster ((ΔOD_620nm_/min)/µM; p < 0.001) and to a higher extent (ΔOD_620nm_; p = 0.066) (**Figures 1B and C**). These findings were complemented with MIC determination, following the EUCAST guidelines, which showed that MICs for cpl-1 were significantly lower than those observed for cpl-7s (**Figure 1D**). Overall, it became evident that cpl-1 consistently outperformed cpl-7s, leading to the decision to proceed exclusively with the cpl-1 endolysin in subsequent assays.

**Figure 1.**
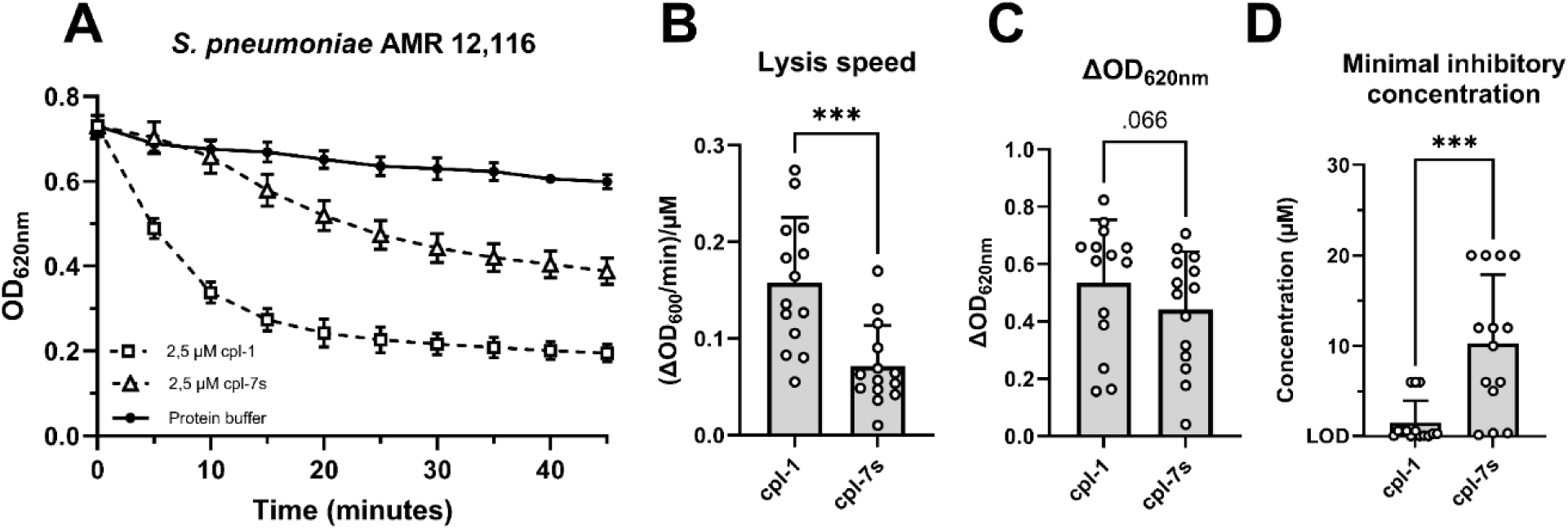
Antibacterial activity of cpl-1 and cpl-7s endolysins against antibiotic-resistant pneumococcal clinical isolates. (**A**) cpl-1 and cpl-7s endolysins at 2.5 µM concentration displayed a strong antimicrobial activity towards the pneumococcal isolate AMR 12,116 in turbidity reduction assays. (**B and C**) Lysis speed (B) and ΔOD_620nm_ (C) analysis showed that cpl-1 hydrolyzed the bacterial peptidoglycan of all the isolates significantly faster and to a higher extent than cpl-7s. (**D**) Determination of minimal inhibitory concentrations (MICs) further confirmed that MICs for cpl-1 were significantly (p < 0.001) lower than those observed for cpl-7s for all pneumococcal clinical isolates tested. In A, datapoints show the mean ± standard deviation of three biological replicates every 5 minutes. In B and C, each dot represents one pneumococcal clinical isolate (14 in total), and bars show the mean ± standard deviation; Lysis speed and ΔOD_620nm_ were analyzed by means of a paired two-tailed t-test, whereas MICs were analyzed by a paired, two-tailed Wilcoxon test (MIC data are not normally distributed); LOD indicates the limit of detection (0.04 µM); * indicates p < 0.05, and *** indicates p < 0.001.

### The cpl-1 endolysin retains its bacteriolytic activity against antibiotic-resistant pneumococcal isolates in human blood and cerebrospinal fluid

To mimic the human pathophysiological conditions during bacteremia, sepsis and meningitis, the activity of the cpl-1 endolysin was evaluated against the penicillin-resistant isolate AMR 19,000, and the multidrug penicillin– and erythromycin-resistant isolates AMR 12,116 and AMR 68,715 in blood obtained from healthy volunteers (blood groups A-, A+ and B+) and in pooled human cerebrospinal fluid. Cpl-1 endolysin significantly reduced the bacterial load in human blood by 1.83 ± 0.61 log_10_ CFU/mL (p < 0.05) for AMR 19,000, and to the limit of detection (i.e., 200 CFU/mL) by 2.52 ± 0.28 and 3.04 ± 0.11 log_10_ CFU/mL (p < 0.001 for both) for AMR 12,116 and AMR 68,715, respectively (**Figure 2A**). In pooled human cerebrospinal fluid, the lytic activity of the cpl-1 endolysin was retained as peptidoglycan hydrolysis was observed by means of TRA (**Figure 2B**).

**Figure 2.**
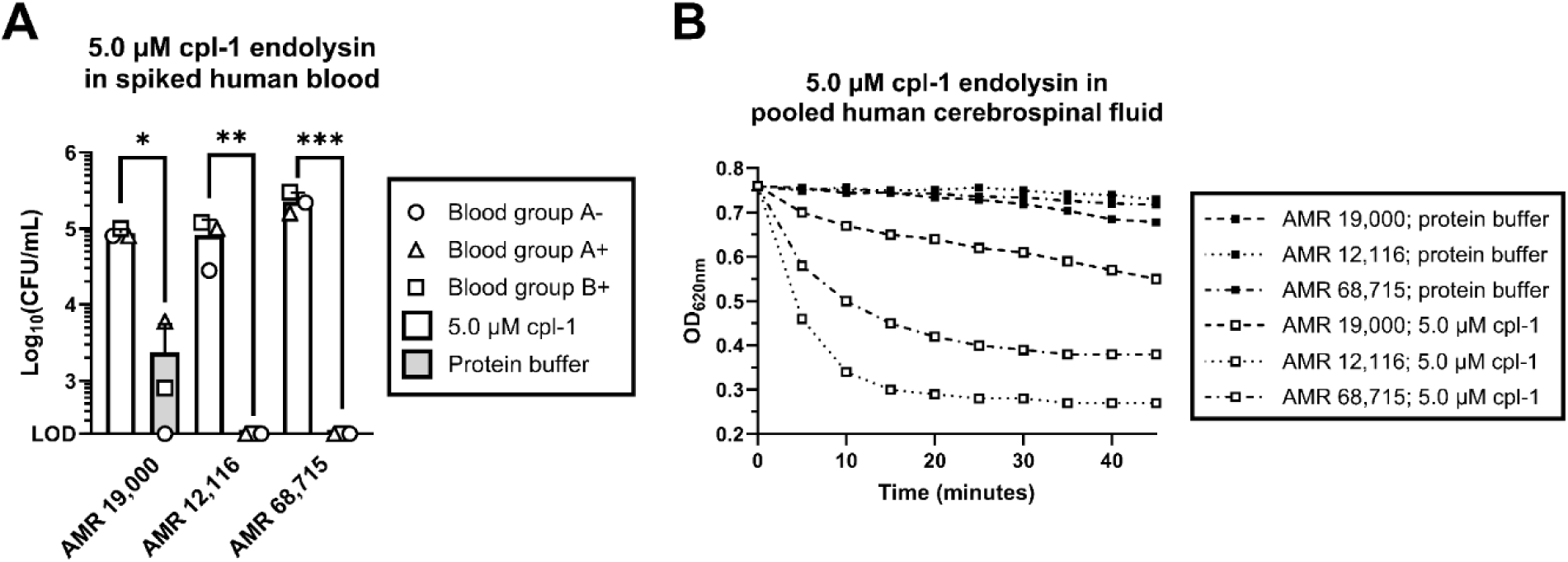
Cpl-1 endolysin retains its antibacterial activity against antibiotic-resistant pneumococcal isolates in human blood and human cerebrospinal fluid. (**A**) Blood obtained from healthy volunteers was spiked with penicillin-resistant AMR 19,000, or penicillin– and erythromycin-resistant AMR 12,116 and AMR 68,715, which was subsequently treated with 5.0 µM endolysin cpl-1 or protein buffer as a negative control. (**B**) Turbidity reduction assay with the same isolates in pooled human cerebrospinal fluid. In A, bars show the mean ± standard deviation of three biological replicates. In B, datapoints show one biological replicate every 5 minutes. Log_10_(CFU/mL) were analyzed by means of a two-tailed, paired t-test; LOD indicates the limit of detection (200 CFU/mL); * indicates p < 0.05, ** indicates p < 0.01 and *** indicates p < 0.001.

### The cpl-1 endolysin lowers the MIC of antibiotic-resistant isolates below the clinical breakpoints by synergizing with penicillin, and through additive effects with erythromycin

To evaluate the impact of endolysin treatment as adjunct to standard-of-care antibiotics in the case of antibiotic resistance, checkerboard assays were conducted to assess potential synergistic, additive, or antagonistic interactions between penicillin, erythromycin, and the cpl-1 endolysin. The fractional inhibitory concentration (FIC) calculated from these checkerboard assays was consistently ≤ 0.50 for penicillin and endolysin cpl-1, which indicates a synergistic effect of cpl-1 towards penicillin (**Table 2**). Even though synergy was not observed for erythromycin and endolysin cpl-1, the FIC values were between 0.50 and 4.00, still indicating that important additive effects of cpl-1 are present towards erythromycin (**Table 3**). A remarkable finding for both antibiotics was that supplementation with endolysin cpl-1 substantially lowered the MIC values of the antibiotics very frequently below the clinical breakpoint for resistance (**Tables 2 and 3**). Of great clinical importance, the MIC values for penicillin (4.00, 4.00 and 16.00 µg/mL) were lowered 8-, 4– and 16-fold (to 0.50, 1.00 and 1.00 µg/mL) for AMR 12,116, AMR 19,000 and AMR 68,715, respectively, and thus all were consistently below the 2.00 µg/mL breakpoint (**Table 2**). For erythromycin, MIC values (4.00, 0.50 and 0.50 µg/mL) were lowered 8-, 8– and 32-fold (to 0.50, 0.06 and 0.01 µg/mL) for AMR 6A, AMR 12,116 and AMR 68,715, respectively, and were below the 0.25 µg/mL breakpoint for the two latter isolates (**Table 3**). This strongly suggests that cpl-1 could serve as a supplemental therapy alongside current standard-of-care antibiotics, potentially resensitizing pneumococci that have developed antibiotic resistance.

**Table 2.** Checkerboard assays between penicillin and the cpl-1 endolysin with the penicillin-resistant pneumococcal isolates. The table indicates minimal inhibitory concentrations observed for penicillin, endolysin cpl-1 or the different combinations thereof. FIC is the fractional inhibitory concentration, which indicates synergy if values are ≤ 0.50. The EUCAST breakpoint for penicillin resistance is set at 2.00 µg/mL.

|  | AMR 12,116 | AMR 19,000 | AMR 68,715 |
| --- | --- | --- | --- |
| Cpl-1 | 5.00 µM | 2.50 µM | 2.50 µM |
| Penicillin | 4.00 µg/mL | 4.00 µg/mL | 16.00 µg/mL |
| Penicillin + cpl-1 | 0.50 µg/mL | 1.00 µg/mL | 1.00 µg/mL |
| Cpl-1 + penicillin | 0.62 µM | 0.62 µM | 0.62 µM |
| FIC | 0.25 | 0.50 | 0.31 |

**Table 3.**
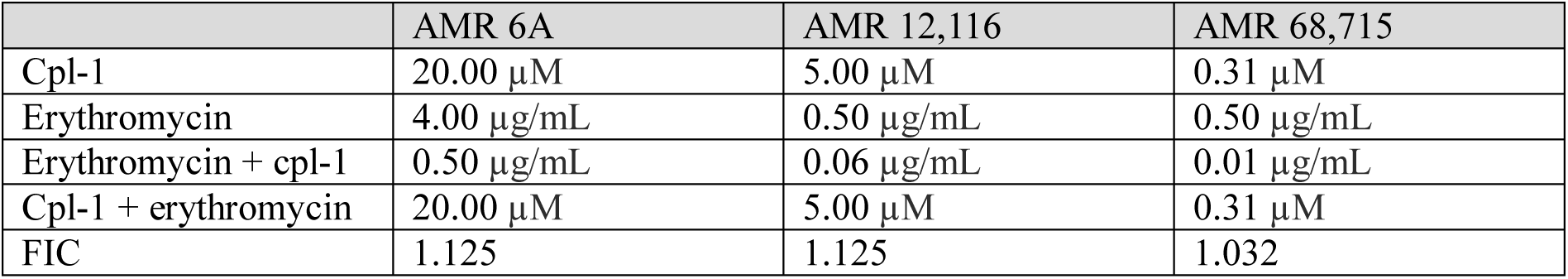
Checkerboard assays between erythromycin and the cpl-1 endolysin with the erythromycin-resistant pneumococcal isolates. The table indicates minimal inhibitory concentrations observed for erythromycin, endolysin cpl-1 or the different combinations thereof. FIC is the fractional inhibitory concentration, which indicates additive effects if values are between 0.50 and 4.00. The EUCAST breakpoint for erythromycin resistance is set at 0.25 µg/mL.

|  | AMR 6A | AMR 12,116 | AMR 68,715 |
| --- | --- | --- | --- |
| Cpl-1 | 20.00 µM | 5.00 µM | 0.31 µM |
| Erythromycin | 4.00 µg/mL | 0.50 µg/mL | 0.50 µg/mL |
| Erythromycin + cpl-1 | 0.50 µg/mL | 0.06 µg/mL | 0.01 µg/mL |
| Cpl-1 + erythromycin | 20.00 µM | 5.00 µM | 0.31 µM |
| FIC | 1.125 | 1.125 | 1.032 |

### Supplementation of cpl-1 endolysin to penicillin or erythromycin treatment reduces bacterial growth and mitigates cytotoxicity in human neuronal cells infected with antibiotic-resistant pneumococcal isolates

To provide preclinical proof-of-concept, human neuronal cells were infected with the penicillin-or erythromycin-resistant pneumococcal clinical isolates and subjected to various treatments that were initiated 90 minutes post-infection, while monitoring bacterial growth and neuronal cell death (neuronal cytotoxicity) up to 6 hours. These treatments included: (i) penicillin or erythromycin at their clinical resistance breakpoints 2.00 µg/mL and 0.25 µg/mL, respectively, (ii) a single dosage of 2.5 µM cpl-1 endolysin, (iii) three serial dosages of 2.5 µM cpl-1 endolysin, or (iv) an “antibiotic + endolysin” combination strategy following the same dosage. An untreated control was included for comparison.

As expected, under penicillin treatment, a similar increase of bacterial growth and neuronal cytotoxicity was observed to that of the untreated control across all isolates due to penicillin resistance (**Figure 3A**). It was striking that, in the case of the AMR 68,715 isolate, a single dose of endolysin successfully eliminated all pneumococci and prevented the rise of neuronal cell death, making multiple dosing or an antibiotic combination strategy unnecessary for this isolate specifically (**Figure 3A**). The clinical isolates AMR 12,116 and AMR 19,000 were less susceptible to the cpl-1 endolysin, with a single cpl-1 dose leading to an increase of bacterial regrowth and neuronal cytotoxicity, especially for AMR 19,000. Nevertheless, administering endolysin three consecutive times still showed a discrete improvement compared to a single dose. Remarkably, for all penicillin-resistant isolates, combining cpl-1 endolysin with penicillin, exploiting their previously identified synergy, resulted in a dramatic reduction of bacterial growth and nearly completely mitigated neuronal cytotoxicity (**Figure 3A**).

**Figure 3.**
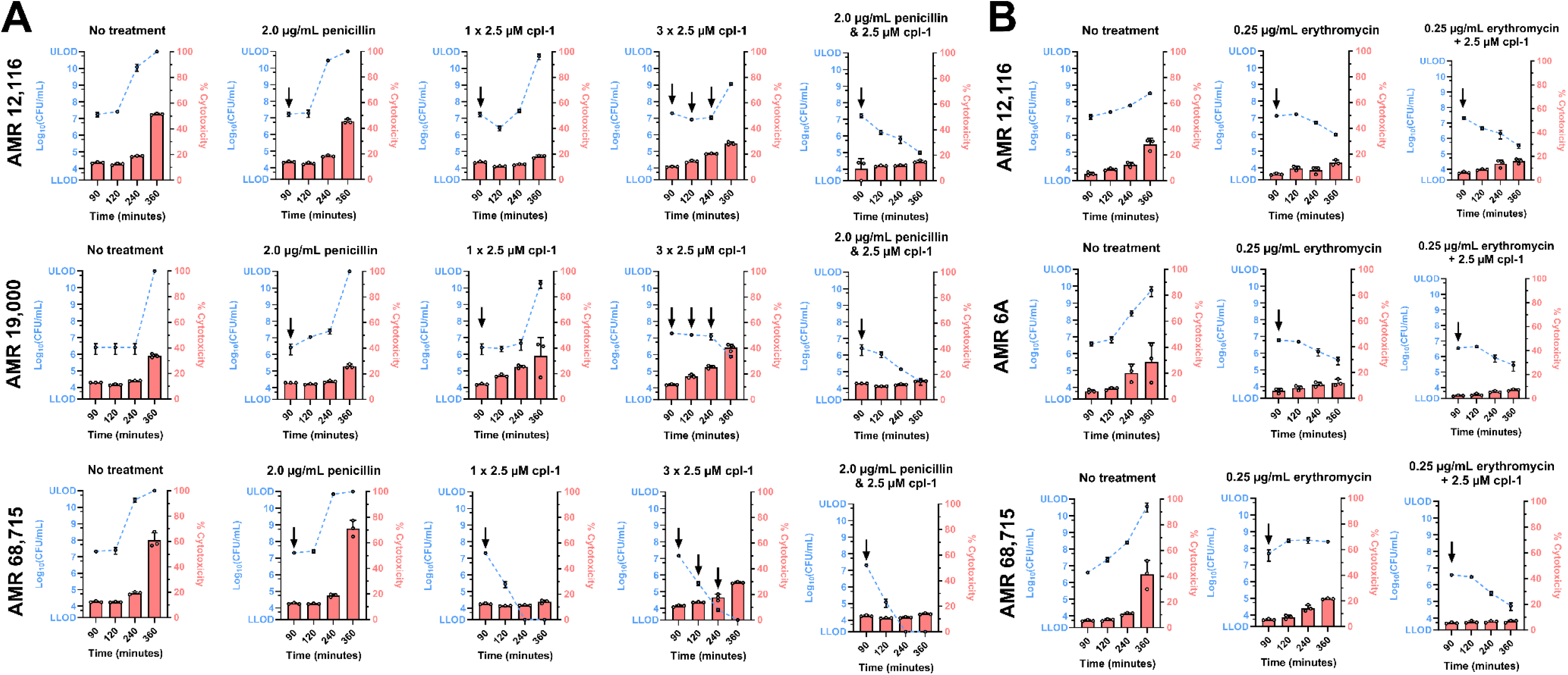
Supplementation of cpl-1 endolysin to penicillin or erythromycin protects human neuronal cells from cytotoxic effect of penicillin– and erythromycin-resistant pneumococcal infections. (**A**) Penicillin and cpl-1 endolysin treatments were evaluated separately, including a multiple dosing strategy for the cpl-1 endolysin, and together as a combination in human neuronal cells infected with the penicillin-resistant pneumococcal clinical isolates AMR 12,116, AMR 19,000, and AMR 68,715. (**B**) Erythromycin treatment was evaluated as a stand-alone treatment, and together with endolysin cpl-1 as a combination, in human neuronal cells infected with the erythromycin-resistant pneumococcal clinical isolates AMR 12,116, AMR 6A, and AMR 68,715. In both A and B, untreated controls were included for comparison; the blue dotted lines represent bacterial growth in log_10_(CFU/mL) plotted on the left y-axis, with data points at 90-, 120-, 240-, and 360-minutes post-infection showing the mean ± standard deviation of three biological replicates. The red bars depict the percentage of cytotoxicity in human neuronal cells at these same time points, plotted on the right y-axis, showing three biological replicates (corresponding to three white circles), alongside bars indicating the mean and an error bar indicating the standard deviation. 2.0 and 0.25 µg/mL are the clinical breakpoints for penicillin and erythromycin resistance, respectively. The black arrow indicates administration of treatment. LLOD and ULOD indicate the lower and upper limit of detection of 200 and 10^11^ CFU/mL, respectively.

In the case of erythromycin treatment, bacterial growth did not increase to the same extent as in the untreated controls, despite the isolates being previously identified as erythromycin-resistant through MIC determination. Nevertheless, the combination of cpl-1 endolysin and erythromycin was found to be the most effective in reducing bacterial growth and further decreasing neuronal cytotoxicity (**Figure 3B**).

Overall, while endolysins alone may not always be effective, even when administered in multiple doses, due to their short half-life and the capability of the surviving bacteria to regrow^17,26^, the addition of cpl-1 to a standard-of-care antibiotic appears to be a very promising therapeutic strategy against antibiotic-resistant bacterial infections, especially in case of meningitis due to the great capability of cpl-1 to protect neurons and prevent neuronal cell death.

### Supplementation of cpl-1 endolysin to penicillin successfully treats penicillin-resistant invasive pneumococcal disease *in vivo*

We then decided to further evaluate the synergistic combination of cpl-1 endolysin with penicillin in our established bacteremia-derived meningitis mouse model alongside placebo (i.e., PBS), endolysin stand-alone and antibiotic stand-alone treatment. Notably, the AMR 12,116 pneumococcal isolate stand out as particularly interesting given that Iovino et al. previously demonstrated that the piliated 35B serotypes usually belongs to the ST-558 pathotype and is a frequent cause of IPD in humans^33^. Furthermore, the 35B serotype is not included in the current pneumococcal vaccines and frequently has reduced penicillin susceptibility^34^.

Within 13 h, all infected mice in the placebo group developed symptoms indicative of IPD and reached experimental endpoints (**Figures 4A and 4B**). Similarly, all penicillin-treated mice succumbed within 21 h, also reaching experimental endpoint of disease (**Figures 4A and 4B**). In contrast, mice treated with cpl-1 endolysin showed a slightly improved probability of survival, most likely due to the antibacterial activity of cpl-1 towards penicillin-resistant pneumococcal infection, which was not affected by the penicillin treatment (**Figures 4A and 4B**); however, due to the short half-life of endolysin, the antibacterial effect was only temporary, and, although the survival of this group was longer than animals treated with penicillin, most of the mice still reached experimental endpoint within 27 h (**Figures 4A and 4B**). On the other hand, all mice treated with the combination strategy remained asymptomatic throughout the first 21hrs, displaying a significantly increased probability of survival compared to the other groups, achieving 100% survival up to 25 h post-treatment, and 89% survival at the end of the experiment. Notably, this reduced survival rate was due to only one mouse developing symptoms at the very end of the experiment (**Figures 4A and 4B**).

**Figure 4.**
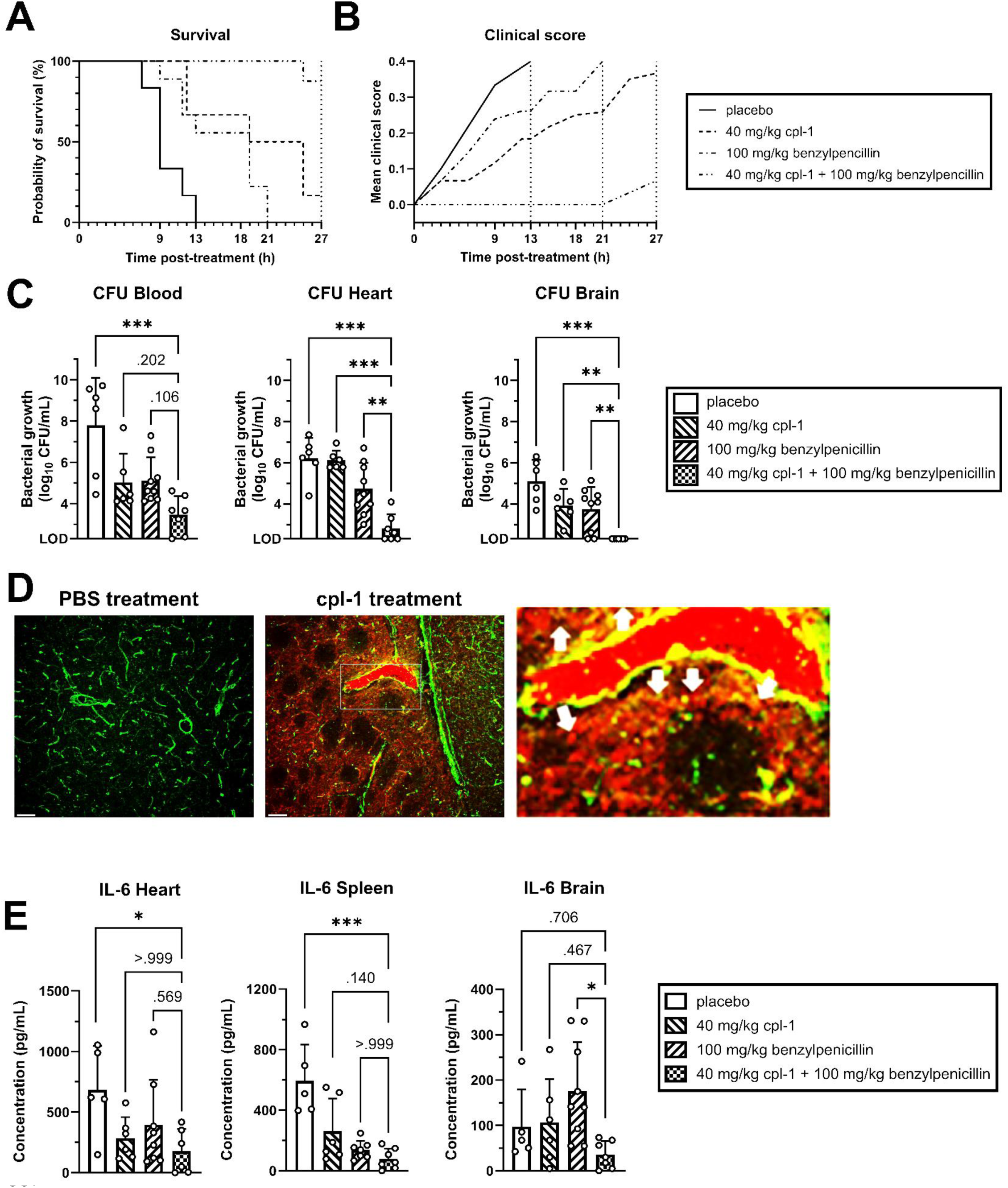
Administration of cpl-1 endolysin supplemented to penicillin protects mice from penicillin-resistant invasive pneumococcal disease. Kaplan–Meier survival curve (**A**) and mean clinical scores (**B**), assessed according to Karolinska Institutet’s veterinary protocol for rodent physiological and psychological well-being. A clinical score of 0.0 indicates a fully healthy status, while 0.4 indicates the humane endpoint. (**C**) Bacterial load in the blood, heart, and brain. (**D**) Confocal images display representative merged z-stacks of three brain tissue sections from two different mice that were stained for brain microvasculature (green) and cpl-1 endolysin (red); brain tissue area within white borders was digitally enlarged 10X to highlight the diffusion (white arrows) of the cpl-1 red signal from the vasculature into the brain parenchyma. The scale bars represent 50 µm. (**E**) Levels of IL-6 in the heart, spleen, and brain. In C and E, data were analyzed using one-way ANOVA with Bonferroni post-hoc tests after outlier removal via the ROUT method (Q = 5%); Bars represent the mean ± standard deviation of biological replicates (each dot represents one mouse); LOD indicates the limit of detection (200 CFU/mL); * indicates p < 0.05, ** indicates p < 0.01 and *** indicates p < 0.001.

The group that received the combination treatment had the highest reduction of bacterial load in the blood, heart and spleen compared to all other groups (**Figure 4C** and **Supplementary Figure S2**); remarkably, the combination treatment completely eliminated bacteria in the brain (**Figure 4C**). Given the remarkable observation that no pneumococci were recovered from the brains of mice that were treated with the combination therapy, we hypothesized that cpl-1 endolysin can cross the blood-brain barrier (BBB) efficiently. In fact, confocal microscopy analysis of brain sections from mice intravenously treated with cpl-1 clearly showed the presence of cpl-1 inside the brain microvasculature accompanied by a diffuse cpl-1 fluorescent signal corresponding to the endolysin in the brain parenchyma, which was absent in the brain of PBS-injected mice (**Figure 4D and Supplementary Figure S3**), therefore demonstrating the efficient capability of cpl-1 to cross the BBB upon systemic administration.

Along with the significant reduction of bacterial load, the impact of antimicrobial treatment on systemic and neuroinflammation was evaluated by quantifying levels of IL-6, an inflammatory cytokine often used as clinical marker in patients with bacterial sepsis^35–37^, in the heart, spleen and brain. Remarkably, IL-6 levels were the lowest for the combination treatment in the heart, spleen, and brain compared to all other groups (**Figure 4E**). This last finding finally showed that the supplementation of cpl-1 endolysin to penicillin dramatically dampens acute inflammation during systemic antibiotic-resistant infection.

## Discussion

IPD is without doubt a major cause of morbidity and mortality globally^38^. *S. pneumoniae* is also the leading cause of bacterial meningitis worldwide, mainly affecting children, with high risk of mortality or long-term neurological sequelae upon survival^5–7,39^. Bacterial meningitis is treated with antibiotics, but even β-lactams, which are frequently used for streptococcal infections, have very low capability to penetrate the BBB and reaching the brain^40^. Furthermore, even in case of BBB penetration, antibiotic-resistance is a constant threat to face in clinics^41^. Moreover, the rise of β-lactam– and macrolide-resistant strains is worrying^10–12^. Endolysins, such as cpl-1 and cpl-7s, have been previously characterized for their antibacterial activity ^25,26^, but fundamental proofs of their efficacy using human-derived tissue, human cells and animal models for meningitis were still missing. In this study, we delivered important preclinical proof-of-concept evidence regarding the therapeutic potential of the cpl-1 endolysin for treating invasive pneumococcal disease and meningitis, protecting human neuronal cells from the cytotoxic effect of the infection, and dampening inflammation, thanks to its capability of restoring antibiotic sensitivity in antibiotic-resistant infections.

Firstly, we demonstrated that all the antibiotic-resistant pneumococcal clinical isolates used in this study were sensitive to endolysin treatment in both human blood as well as human cerebrospinal fluid, an observation with huge clinical importance that has not yet been reported. We further characterized the impact of cpl-1 endolysin supplementation to penicillin or erythromycin treatments in resistant pneumococcal strains. Notably, our results revealed that cpl-1 supplementation significantly reduced the MIC of penicillin– and erythromycin-resistant isolates. Moreover, we are the first to demonstrate that this reduction lowers MIC values below clinical resistance breakpoints in a standardized setting as outlined by ESCMID guidelines^27^. We found this interaction to be synergistic with penicillin, and additive with erythromycin. This marked difference between penicillin and erythromycin can most likely be explained by their different methods of action. Whereas erythromycin blocks the bacterial protein synthesis, penicillin and endolysins both target the pneumococcal cell wall, thus potentially levering each other^14,15,42^. Following this observation, our findings strongly suggest that endolysins may be primarily used synergistically with cell wall-targeting antibiotics to restore antibiotic susceptibility^14,15^, in line with cell membrane-targeting antibiotics^23^. Furthermore, this reduction in MIC also has important pharmacokinetic implications, given that the effectiveness of antimicrobial treatment *in vivo* primarily depends on the duration that drug concentrations remain above the MIC (i.e., T_>MIC_)^43^. The observed synergistic effects between penicillin and cpl-1 endolysin suggest that lowering the MIC of both agents can substantially extend the T_>MIC_ during antimicrobial treatment. Based on previously reported half-lives of benzylpenicillin and the cpl-1 endolysin in experimental IPD models^26,43^, we calculated that penicillin treatment alone would fail after 3 h in our bacteremia-derived meningitis mouse model, whereas supplementing penicillin treatment with cpl-1 endolysin should extend the T_>MIC_ as far as 25 h. Consequently, all mice treated with penicillin alone rapidly developed symptoms and succumbed within 21 h, whereas those receiving the combination therapy indeed survived significantly longer.

Another critical gap that we addressed is whether endolysins should be clinically implemented as a stand-alone treatment, or in combination with standard-of-care antibiotics^17,44^. Both our *in vitro* and *in vivo* experiments clearly demonstrate that endolysin treatment, thus without antibiotic supplementation, could provide therapeutic relief in the case of antibiotic-resistant infections thanks to its strong antibacterial activity; however, due to the short half-life of endolysin, as previously reported^17,26^, pneumococci surviving the endolysin treatment could regrow again. In marked contrast, the combination of endolysin cpl-1 with either penicillin or erythromycin consistently rescued human neuronal cells from antibiotic-resistant infection after only one dosage. We importantly confirmed these findings *in vivo*; in fact, a single cpl-1 endolysin dose together with penicillin significantly reduced penicillin-resistant pneumococci in the blood, heart and spleen, and, remarkably, completely eliminated bacterial presence in the brain. Consequently, mice receiving this combination strategy had a remarkable 89% survival and remained asymptomatic over the course of the infection experiment, with a slight increase of clinical score in the last hours of the experiment due to only one mouse in that group that developed symptoms. The improved survival, symptomatology and bacterial load in the tissues were also confirmed by the reduced inflammation observed in mice that received the combined treatment compared to the groups that received either the endolysin or penicillin alone. Thus, our data strongly suggests that if clinical implementation of endolysin is aimed, which is expected according to the WHO^13,17^, their strength lies in a combinatorial approach with antibiotics, irrespective of the resistance levels of the targeted bacteria to the combined antibiotic. This important concept has not previously been reported in the field of Infectious Diseases.

Finally, meningitis is today a dramatic burden in Public Health because of the permanent neurological disabilities caused by neuronal damage inflicted by bacteria in the brain^6,7^. We here provide for the first time clear evidence that endolysin cpl-1 is well capable of penetrating the BBB and successfully reaching the brain upon systemic administration. Together with our observation of human neuronal cells that were successfully rescued from the cytotoxic effect of antibiotic-resistant infection, our findings highlight the incredible potential that cpl-1 has to prevent neurological impairment during meningitis onset, achieving what antibiotics cannot do due to their low capability to penetrate the BBB and due to the increasing emergence of antibiotic resistance^40^.

In conclusion, this work demonstrates that pneumococci resistant to penicillin and erythromycin can be resensitized to antibiotic treatment through endolysin supplementation. This approach lowers MIC values below clinical resistance breakpoints via synergistic or additive effects with penicillin or erythromycin, respectively. *In vivo*, this re-sensitization leads to significant bacterial load reductions systemically, and to the complete clearance of bacterial presence in the brain; a strong antibacterial effect that is accompanied by a reduced systemic and neuroinflammation and that, overall, contributes to an asymptomatic course of infection and a significantly increased probability of survival. Moreover, the ability of endolysins to effectively cross the BBB, protect human neuronal cells from the cytotoxic effects of antibiotic-resistant infections, and completely eradicate antibiotic-resistant pneumococci in the brain, highlights its remarkable potential as an adjunct to antibiotic treatment not only for invasive diseases, such as bacteremia, but also for pneumococcal meningitis.

## Acknowledgements

We would like to acknowledge Prof. Anders P. Håkansson at Lund University and the Culture Collection University of Gothenburg (CCUG) for providing the pneumococcal isolates used in this work. We would also like to acknowledge the technical assistance of Dr. Miguel T. Vian and Ida Karimpour.

## Author contributions

Conception or design of the work: NV and FI; data collection: NV, LK and KF; data analysis and interpretation: NV, LK, KF and FI; drafting the article: NV and FI; critical revision of the article: NV, LK, KF and FI. All authors have read and agreed to the published version of the manuscript.

## Funding

This research was funded by a postdoctoral scholarship for NV and FI from the Kronprinsessan Lovisas förening för barnasjukvård och Stiftelsen Axel Tielmans minnesfond with grant number 2023-00758, and the grants, holded by FI, from Vetenskapsrådet (The Swedish Research Council), Marcus Bergvall Foundation, Åhlén Foundation, Tysta Skolan Foundation, Wera Ekström Foundation for Pediatric Research, and Gun & Bertil Stohne Foundation.

## Data availability

All data generated or analyzed during this study are included in the published article and are available on the Mendeley Dataset DOI: 10.17632/9b55khv693.1.

## Conflict of interest

NV declares issued and pending patent applications with regards to endolysin technology.

## Notes

https://doi.org/10.17632/9b55khv693.1

### Competing Interest Statement

Niels Vander Elst declares issued and pending patent applications with regards to endolysin technology.

