## Supplementary Figure S1, Supplementary Figure S2A-D, Supplementary Figure S3 for "Bacteriophage-derived endolysins restore antibiotic susceptibility in penicillin- and erythromycin-resistant *Streptococcus pneumoniae* infections"

**Supplementary Figure S1. Comparative analysis of the lytic activity of the cpl-1 and cpl-7s endolysins against the pneumococcal clinical isolates of this study.** cpl-1 and cpl-7s endolysins at 2.5 µM concentration displayed a strong antimicrobial activity towards all pneumococcal isolate of this study in turbidity reduction assays. *S. pneumoniae* AH 16,031 was challenged at a lower $OD_{620nm}$ due to reduced growth *in vitro*. Datapoints show the mean ± standard deviation of three biological replicates every 5 minutes.

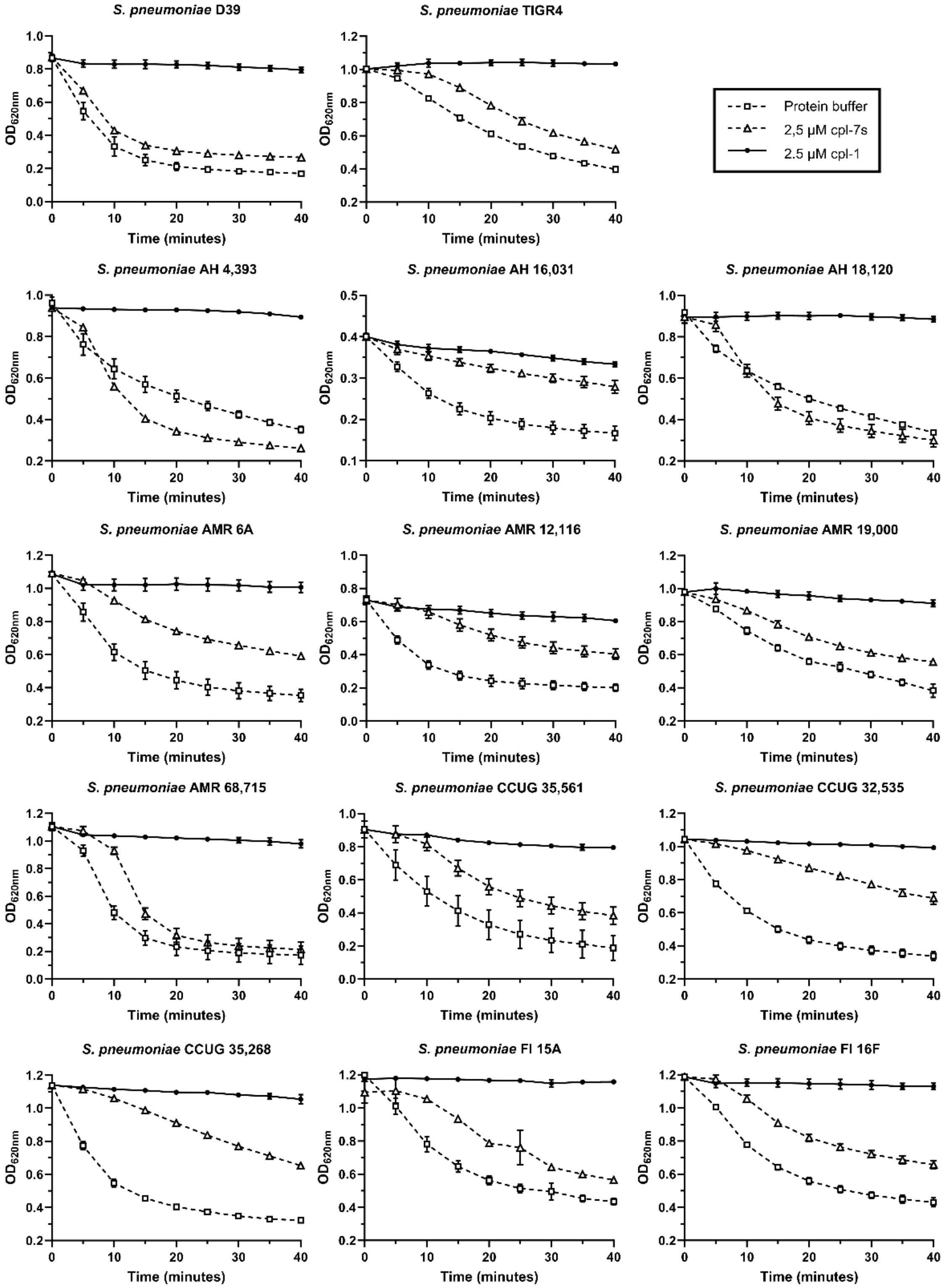

**Supplementary Figure S2. *In vivo* validation of a combination therapy with endolysin cpl-1 and penicillin, compared to stand-alone treatments and placebo (PBS) in a penicillin-resistant bacteremia-derived meningitis mouse model.** Bacterial load in the spleen (**A**). Levels of TNF-α in the heart (**B**), spleen (**C**) and brain (**D**), with one mouse from the PBS group excluded due to consistent outlier values. Data were analyzed using one-way ANOVA with Bonferroni post-hoc tests after outlier removal via the ROUT method (Q = 5%); Bars represent the mean ± standard deviation of biological replicates (each dot represents one mouse); LOD indicates the limit of detection (200 CFU/mL); *** indicates $p < 0.001$.

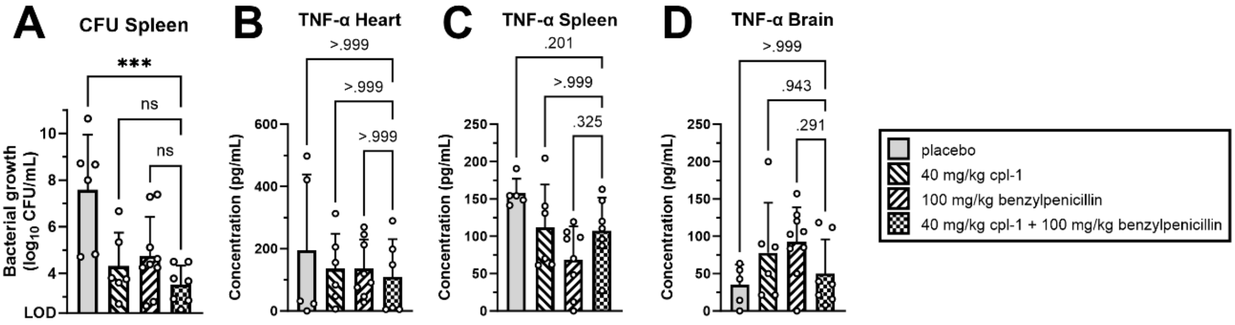

**Supplementary Figure S3. Endolysin cpl-1 efficiently crosses the blood-brain barrier.** Confocal images display representative merged z-stacks (20 images, 20.0 µm thickness) of three brain tissue sections from mice intravenously administered with PBS (n = 2) or 0.75 mg of the cpl-1 endolysin (n = 2). Tissues were immunofluorescent stained for the brain microvasculature (lectin, Alexa Fluor 488, green signal) and the cpl-1 endolysin (hexahistidine-tag, Alexa Fluor 647, red signal). The lower row depicts merged images that combine the staining for the microvasculature with the cpl-1 endolysin showing no cpl-1 signal for the PBS-treated group, as expected, and diffusion of the red signal from the vasculature into the brain parenchyma for the cpl-1-treated group. Scale bars represent 50 µm.

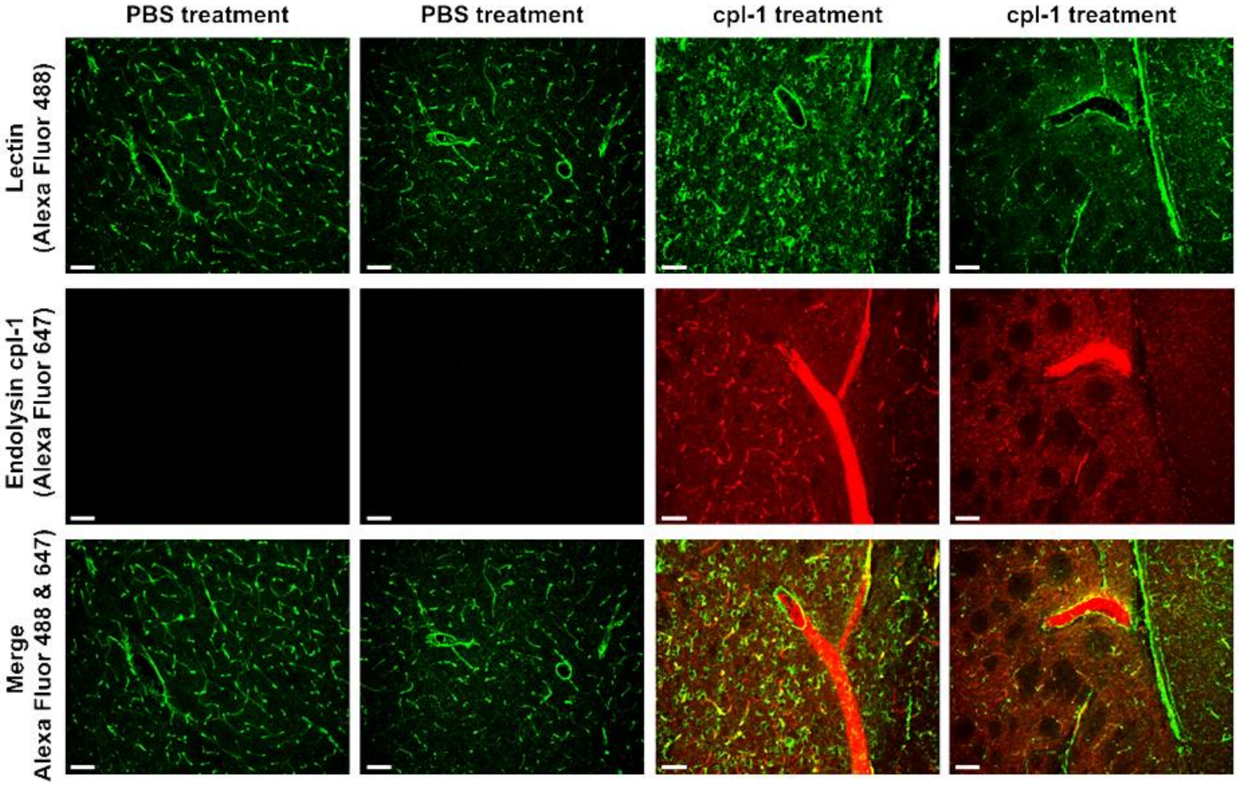
